# Regulating biocondensates within synthetic cells via segregative phase separation

**DOI:** 10.1101/2024.10.18.619037

**Authors:** Chang Chen, Caroline M. Love, Christopher F. Carnahan, Ketan A. Ganar, Atul N. Parikh, Siddharth Deshpande

## Abstract

Living cells orchestrate a myriad of biological reactions within a highly complex and crowded environment. A major factor responsible for such seamless assembly are the preferential interactions between the constituent macromolecules, either associative or segregative, that can drive de-mixing to produce co-existing phases, and thus provide a dynamic intracellular compartmentalization. But how these two types of interactions, occurring simultaneously within the cytoplasmic space, influence each other is still largely unknown. This makes understanding and applying the molecular interactions that interfere with each other in such crowded environments crucial when engineering increasingly complex synthetic cells. Here, we show that the interplay between segregative and associative phase separation within cell-mimicking vesicles can lead to rich dynamics between them. Using on-chip microfluidic systems, we encapsulate the associative and segregative components in cell-sized containers and trigger their phase separations to create hierarchical structures that act as molecular recruiters, membrane targeting agents, and initiators of condensation. The obtained multiphase architecture provides an isolated microenvironment for condensates, restricting their molecular communication as well as diffusive motion, and leading to budding-like behaviour at the lipid membrane. In conclusion, we propose segregative phase separation as a universal condensate regulation strategy in managing molecular distribution, condensate location, as well as membrane interaction. We believe our approach will facilitate controlling the behaviour of membraneless organelles within synthetic cells.

---

The interior of a living cell is an incredibly crowded and dynamic environment packed with a vast array of biopolymers and organelles. In this extraordinarily crowded aqueous space, proteins, polysaccharides, nucleic acids, and other biomolecules constantly deplete and interact, creating a highly active and bustling ecosystem. Bottom-up synthetic biology is an approach that aims to unravel the underlying fundamental principles by constructing self-assembled, cell-mimicking systems from fundamental biomolecules. This makes understanding and applying the molecular interactions that interfere with each other in such crowded environments crucial when building synthetic cells. Commencing with simple models, such as cell-sized vesicles, one can customize and engineer biological systems, leading to increasingly complex intracellular architectures (*1–3*). Both membrane-bound (*4–7*) and membraneless (*8*) modules can be constructed to achieve this, with the latter being more dynamic and gaining increasing attention (*9–11*).

Within the intracellular environment, membraneless organelles (*12*) serve multifaceted functions (*13–15*) and often interact with cell membranes (*2*). As a result, understanding and applying model membraneless organelles is key to engineering more complex synthetic cell systems, achieving basic cellular functions, and move towards specific applications (*16*). Since the groundbreaking discovery of membraneless organelles such as P granules (*17*) and the nucleolus (*18*), the mechanism behind their formation, the phenomenon of liquid-liquid phase separation (LLPS), has garnered increased recognition (*19, 20*). Based on how the molecules are organized within the resulting phases, LLPS can be broadly characterized as either associative phase separation (APS) or segregative phase separation (SPS). APS occurs when there are strong intermolecular forces, such as charge-based attraction between the molecules, causing them to form a condensed liquid phase, commonly referred to as a coacervate or a condensate (*21*). In contrast, SPS is driven in mixtures of polymers with distinct chemical properties, resulting in immiscible, segregated polymer-rich phases, and thereby minimizing contact between them (*22*).

In recent years, there have been active efforts to mimic cellular systems by triggering APS inside synthetic cells leading to dynamic processes, such as reversible condensation (*23*), division (*24*), intracellular trafficking (*25*), biochemical reactions (*26*), and membrane interactions (*27*). In the cellular environment, the functional effectiveness of condensates is tuned by confining them to distinct locations such as the plasma membrane, cytoplasm, or nucleus (*28*). Thus, replicating these behaviors within synthetic confinements would represent a significant stride in the realm of synthetic cell studies. Despite fruitful advancements, achieving the precise localization of coacervates requires cumbersome procedures including the use of additional ingredients in the membrane and condensates, e.g., charged lipids and condensates (*29*), condensates with hydrophobic properties (*30*), or sealing membrane pores with condensates (*31*). Another challenge is the restriction of the coacervate movement, with existing strategies allowing them to move freely across the entire membrane surface (*30*). This non-specific localization may not always be favorable for coacervate functionality, for example, to attain cell polarity or restrict physical contact with other membrane-bound entities. Last but not least, the cross-talk between condensates and the membrane’s inner leaflet still requires further exploration (*32*). The wetting behavior of condensates is primarily being investigated via interactions with the outer membrane leaflet, leading to shape changes (*33*) and coacervate penetration (*34*). Likewise, triggering intracellular coacervation and their interactions with the membrane from the inner leaflet may lead to budding-or even exocytosis-like behavior (*35*). Thus, gaining spatiotemporal control over internal coacervates is important for engineering synthetic cells.

SPS, on the other hand, is considered an important tool for mimicking cellular behavior in a crowded environment. Utilizing common molecular crowders, SPS is widely applied for intracellular compartmentalization via temperature (*36*), or osmolyte (*37*) trigger. The resulting LLPS can influence morphological changes in the membrane including budding (*38*), tube formation (*39*), asymmetric division (*40*) as well as distribution (*41*), and phase separation of membrane lipids (*22*). Using liposomes, prior studies have used SPS to construct polar structures (*42*), or liposomes with multiple distinct compartments (*22*). However, although APS and SPS are both well-established individually, their simultaneous and dynamic occurrence within the cytoplasmic space remains unclear. Therefore, understanding the interaction between APS and SPS and combining their respective advantages is of great significance for the study of synthetic cells.

In this paper, we explore and utilize SPS-APS interplay in microconfinements to regulate various coacervate behaviors including molecular enrichment, membrane-targeting, restricted diffusion, and budding at lipid membranes. Using polyethylene glycol (PEG)/dextran (DEX) as the SPS system and poly-L-lysine (PLL)/adenosine triphosphate (ATP) as the APS system, we first conducted bulk experiments to demonstrate the recruitment of coacervates and the physicochmically restrictive environment offered by the SPS-induced membrane-free confinements (SMCs). Then, utilizing on-chip microfluidic production techniques, we encapsulated the APS and SPS components within water-in-oil-in-water double emulsions as well as liposomes. Based on these two synthetic cell models, we investigated either free or membrane-bound SMCs in regulating coacervates. Osmolarity-triggered SMCs were found to relocate specific coacervate components (PLL), capable of targeted migration to the membrane. Via a pH trigger, we constructed multiphase coacervate-in-SMC architectures that could be either suspended in the lumen or distributed on the membrane in the form of micron-sized buds. When confined to the membrane by SMCs, the coacervates remained largely isolated from each other and showed highly restricted diffusion. Overall, we achieved diverse coacervate regulation strategies in vesicles using APS-SPS-membrane interplay. This universal strategy provides a new path for mimicking membraneless organelles in synthetic cells and further studying the complex interactions between SPS, APS, and the lipid membrane for advancing synthetic cells.

## SPS physicochemically regulates condensate dynamics

We utilized PEG (8 kDa) and DEX (≈10 kDa), two commonly used chemically dissimilar and electrically neutral polymers, as a model system for SPS (Figure 1a). To facilitate subsequent experiments, we first established binodal curves for PEG/DEX using a variation of cloud-point titration for varying conditions of pH, viscosity, and buffer concentrations (see Methods for details, also see Supplementary Figure **??**). Regardless of the varied conditions, the system exhibited consistent binodal curves (Figure 1b), providing us with a reliable compartmentalization system in the subsequent experiments (Figure 2-4). As a model APS system, we used PLL and ATP, an extensively used molecular pair that undergoes complex coacervation via electrostatic attraction. To begin investigating the APS-SPS interactions, we assessed the tendency of PLL and ATP to get sequestered into one of the SPS phases, by calculating their partition coefficients (*K*) for the two phases (see Methods for details). As can be clearly seen from Figure 1c, there is a pronounced increase in the ratio of PLL fluorescence in the DEX-rich phase over that in the PEG-rich phase (*K*_*PLL*_), with the values increasing significantly further with increasing concentrations of PEG or DEX. On the contrary, *K*_*ATP*_ exhibited a value around 1 irrespective of the PEG/DEX concentrations (see Figure 1d), indicating ATP did not show any significant preference for either phase.

**Figure 1:**
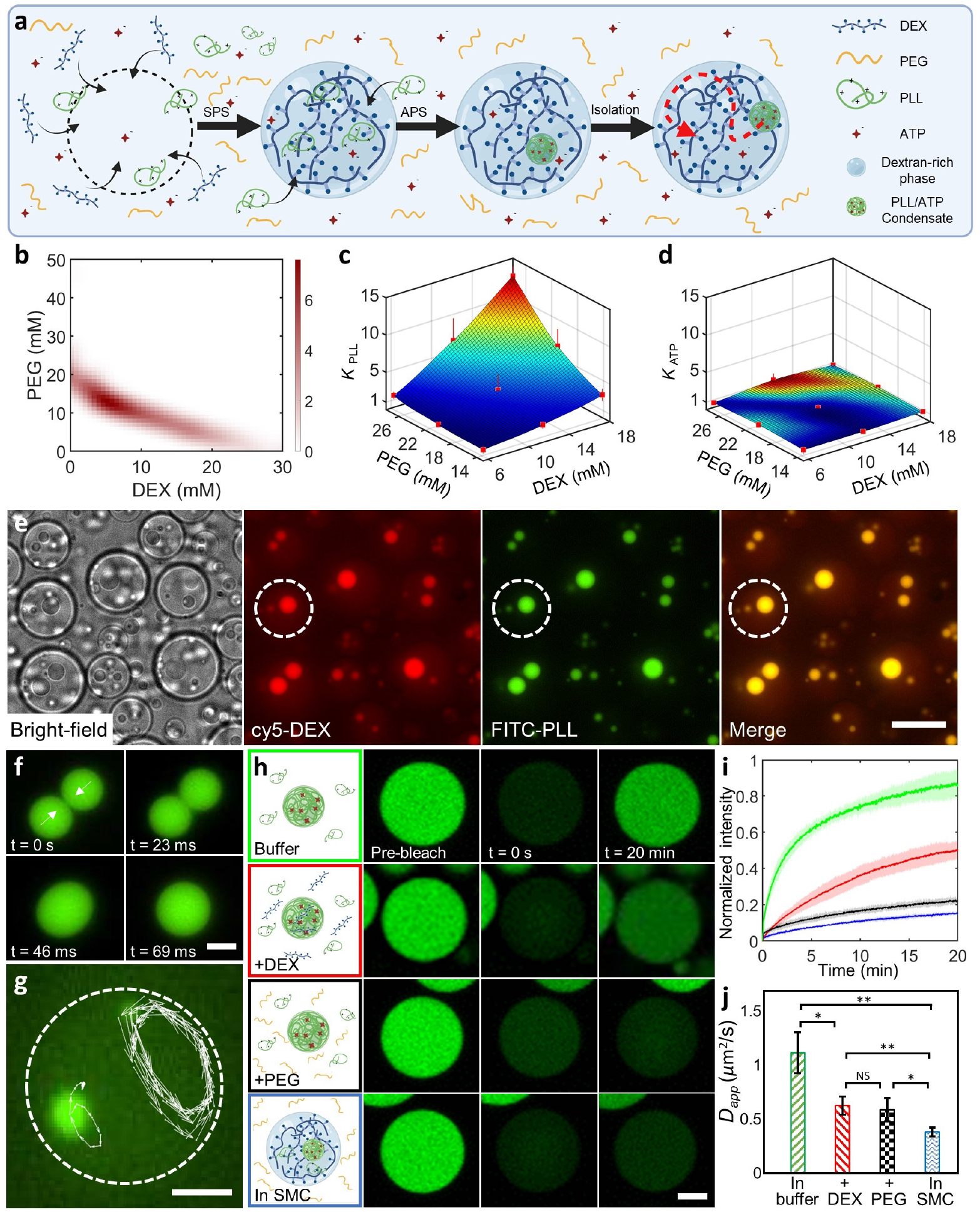
SPS domains regulate APS dynamics. (a) Conceptual sketch showing the SPS-APS interplay: SPS generates a DEX-rich domain that sequesters PLL molecules, inducing PLL/ATP coacervation and restricting coacervate motion within the SMC. (b) A combined binodal curve of PEG (8 kDa) and DEX (10 kDa), obtained by cloud-point titration, under different conditions of pH (3.8 to 9.7), viscosity (with and without glycerol), and ionic strengths (15 and 25 mM Tris-HCl buffer) used throughout this work. Regardless of the conditions, the binodals showed the same trend. The color depth corresponds to the number of data points. (c-d) Three-dimensional surface plots representing the partition coefficients (*K*) of (c) FITC-PLL and (d) cy3-ATP within the SPS system, where K represents the ratio of fluorescence in the DEX-rich phase over that of the PEG-rich phase. Colors represent the extent of partitioning, with red being higher partitioning and blue being lower partitioning. (e) Microscopy images (from left to right: bright-field, DEX fluorescence, PLL fluorescence, merged fluorescence) showing the formation and restriction of coacervates within DEX-rich domains. The dotted circle indicates the boundary of one such SPS domain. As can be seen, fluorescent DEX gets further enriched within the coacervates. Scale bar, 20 μm. (f) PLL/ATP coacervates show their usual liquid-like behavior in the form of droplet fusion within SPS domains. (g) The coacervates remain confined within the DEX-rich domains even in presence of an external fluid flow, as shown by their confined trajectories. Dotted circle indicates the SMC boundary. (h) Time-lapse fluorescence images showing the fluorescence recovery of PLL (3 mg/ml)/ATP (4 mM) coacervates in varying conditions. For f-h, scale bar, 5 μm. (i) Fluorescence recovery curves of FITC-PLL after bleaching the entire PLL/ATP coacervates under varying conditions as indicated in h. (j) Corresponding diffusion coefficients of PLL after fitting the FRAP curves obtained in i. In e-g, the experiments were conducted by mixing 3 mg/ml PLL (unlabelled: FITC-labeled = 5:1, mass ratio), 4 mM ATP, 12 mM DEX and 12 mM PEG. In h-j, the experiments were conducted by mixing 3 mg/ml PLL (unlabelled: FITC-labeled = 5:1, mass ratio) and 4 mM ATP without PEG and DEX (green panel), with 12 mM DEX (red panel), with 12 mM PEG (black panel), and with 12 mM DEX and 12 mM PEG (blue panel). Data points represent mean values derived from three independent measurements, with error bars indicating standard deviations.

**Figure 2:**
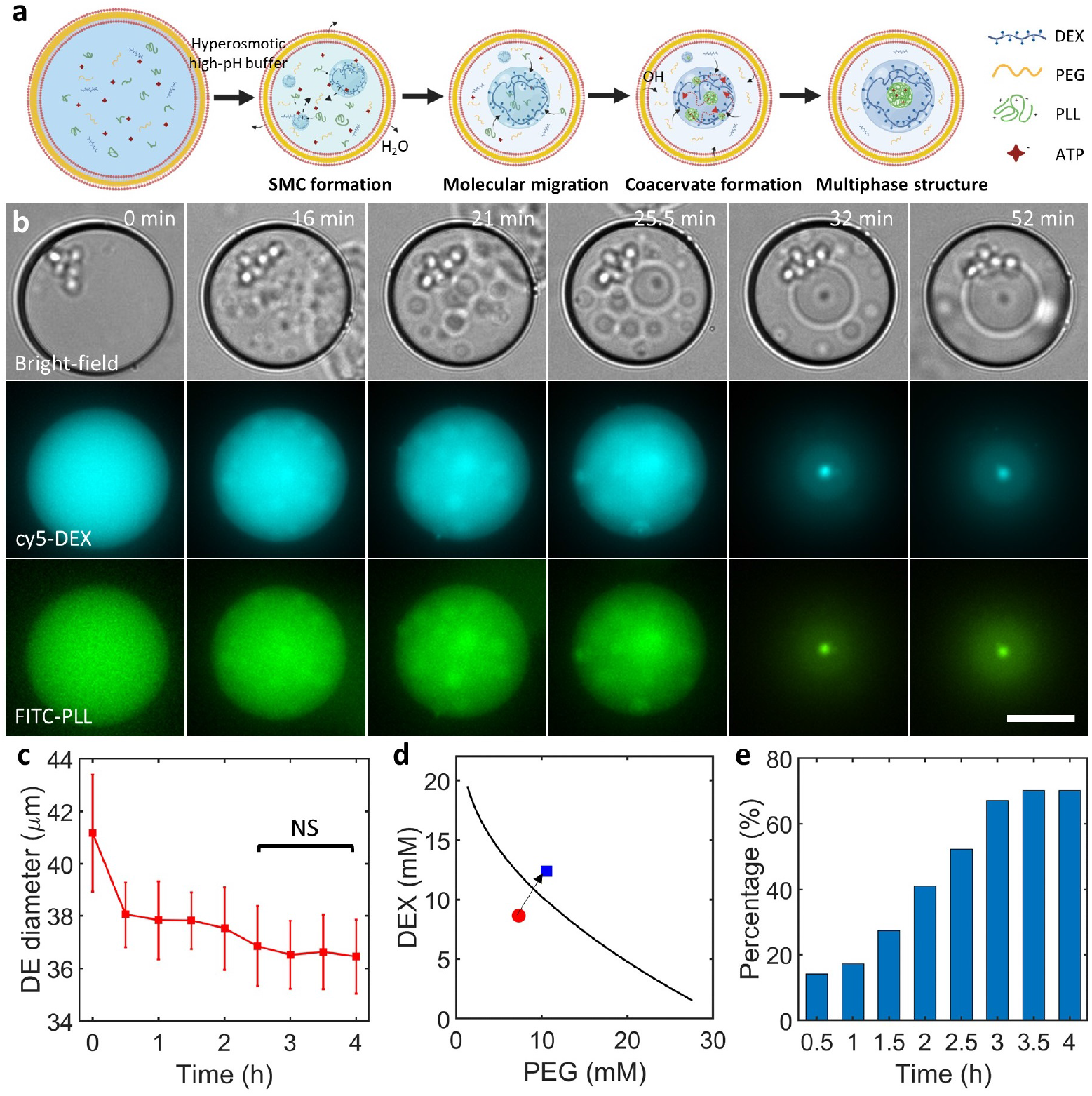
SMC-restricted coacervate dynamics within double emulsion compartments. (a) A conceptual sketch showing coacervate formation and recruitment by SMC, leading to a coacervate-in-SMC structures in double emulsions. (b) Bright-field and fluorescence time-lapse images showing the initial state, coacervate formation, and their restriction within SMCs after feeding the double emulsions with a hypertonic, high-pH buffer. The bright-field images show the double emulsion shrinkage as a result of the hypertonic trigger. The DEX fluorescence (shown in cyan) shows the initiation of SPS, eventually leading to a multiphase structure, with the inner DEX-rich phase, marked by a higher intensity. The PLL fluorescence (shown in green) shows a process almost identical to DEX channel, eventually forming a coacervate within the SMC. The images at 0 min show a representative double emulsion before the trigger. The last two fluorescence panels are taken from the bottom plane while the rest are all captured at the equatorial plane. Scale bar, 20 μm. (c) Corresponding mean diameters of the double emulsion population showing significant reduction in size for the first two hours after the trigger; no significant difference (marked NS) was observed after 2.5 hours. Data points represent mean values while the error bars indicate standard deviations (n ≥ 55 double emulsions for each data point). (d) The initial concentration of the encapsulated PEG and DEX (red circle) crosses the binodal (black curve, obtained by fitting the experimental data) after the trigger (blue square, calculated from the volume change). (e) A plot showing the percentage of double emulsions containing coacervates in SMCs after the trigger (n ≥ 61 double emulsions for each data point). For all the experiments, the encapsulated mixture consisted of 7.3 mM DEX (unlabelled: cy5-labeled = 1000:1, molar ratio), 8.6 mM PEG, 2.4 mg/ml PLL (unlabelled: FITC-labeled = 5:1, mass ratio), 0.8 mM ATP, and 15% v/v glycerol in 15 mM citrate-HCl (pH 4). The double emulsion suspension was combined with an equal volume of feeding aqueous solution containing 500 mM sucrose and 15% v/v glycerol in 15 mM Tris-HCl (pH 9).

**Figure 3:**
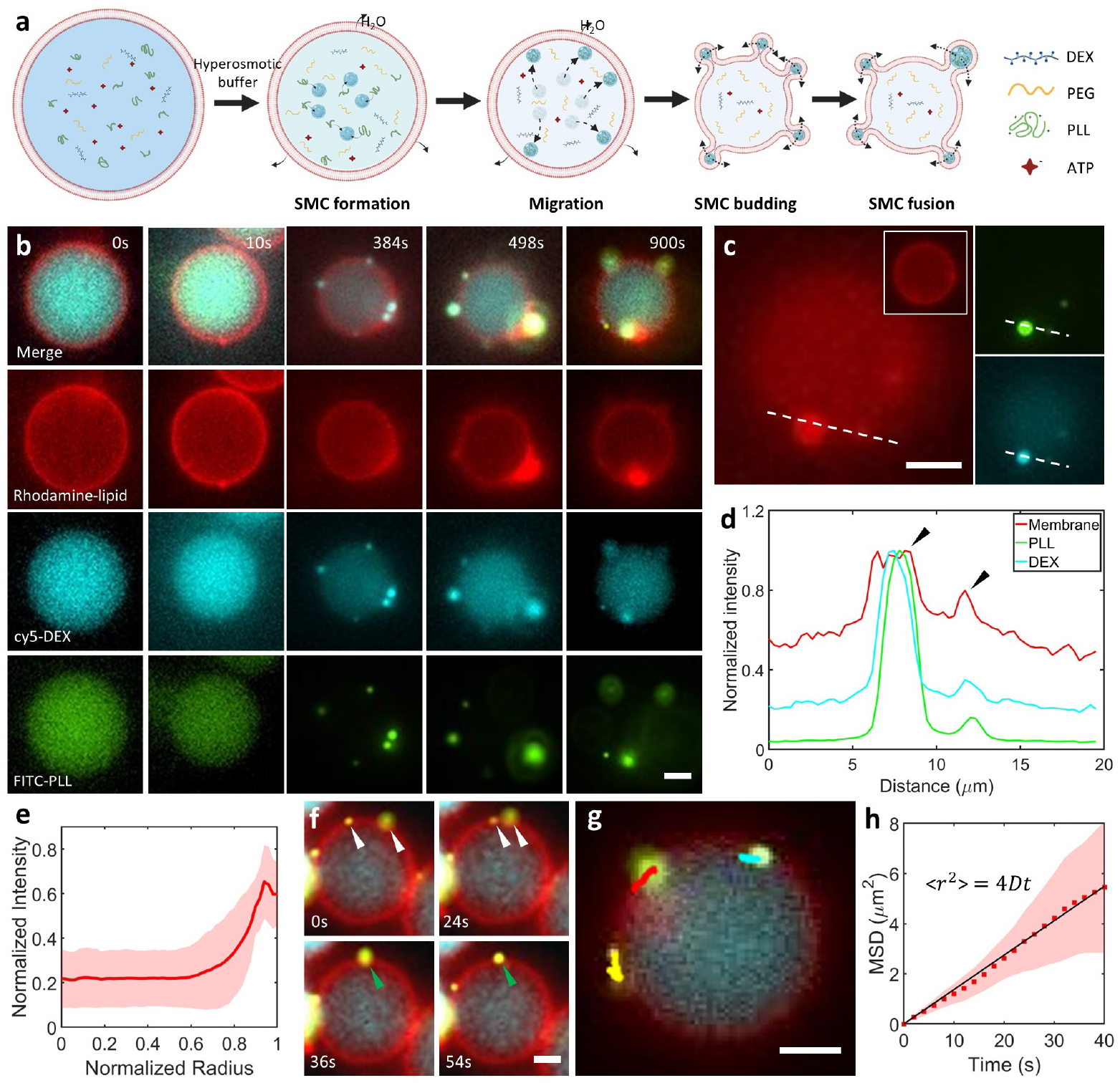
SMCs bud on the membrane and recruit APS components. (**a**) Conceptual sketch showing SMC formation and recruitment of PLL molecules, ultimately resulting in budded structures at the membrane that show a restricted 2D motion and are semi-enclosed by the membrane. (**b**) Fluorescence time-lapse images showing the initial state, followed by SMC formation, PLL recruitment, membrane localization, and budding upon trigger. The budded SMCs can be seen as intense bright dots (in cyan) appearing at the membrane over time. The PLL channel (in green) shows strong sequestration of PLL in the SMCs. The panel at 0 min shows a representative liposome before the trigger. (**c**) Fluorescence image of the liposome surface showing the budded structures; the inset illustrates the equatorial plane, showing the overall spherical nature of the liposome. The images on the right show the PLL (top) and DEX (bottom) fluorescence in the bud. (**d**) Line graphs corresponding to the dotted lines in **c** showing the co-localization of the lipid bud, SMC, and the sequestered PLL molecules. (**e**) Mean PLL fluorescence as a function of the normalized liposome radius (*n* = 10 different liposomes). Shaded area indicates standard deviation. In c-e, the data were collected 30 minutes after the trigger. (**f**) Time-lapse showing a fusion event between two membrane-bound SMCs. (**g**) Trajectories of SMCs over a course of 22 seconds, showing their 2D-restricted movement on the membrane surface. (**h**) A plot showing linear increase in the MSD of the budded coacervates. (*n* = 5 coacervates in 4 different liposomes). The black line shows a linear fit (R^2^ = 0.99). In f-h, the data were collected 20 minutes after the trigger. For all experiments, the lipid composition was 99.9 % DOPC + 0.1% Liss Rhodamine PE (molar ratio). Encapsulating mixture consisted of 7.3 mM DEX (unlabelled: cy5-labeled = 1000:1, molar ratio), 8.6 mM PEG, 2.4 mg/ml PLL (unlabelled: FITC-labeled = 5:1, mass ratio), 0.8 mM ATP, and 15% v/v glycerol in 25 mM citrate-HCl (pH 4). The liposome suspension was combined with an equal volume of feeding aqueous solution containing 600 mM sucrose and 15% v/v glycerol in 25 mM Tris-HCl (pH 9). All scale bar, 5 μm.

**Figure 4:**
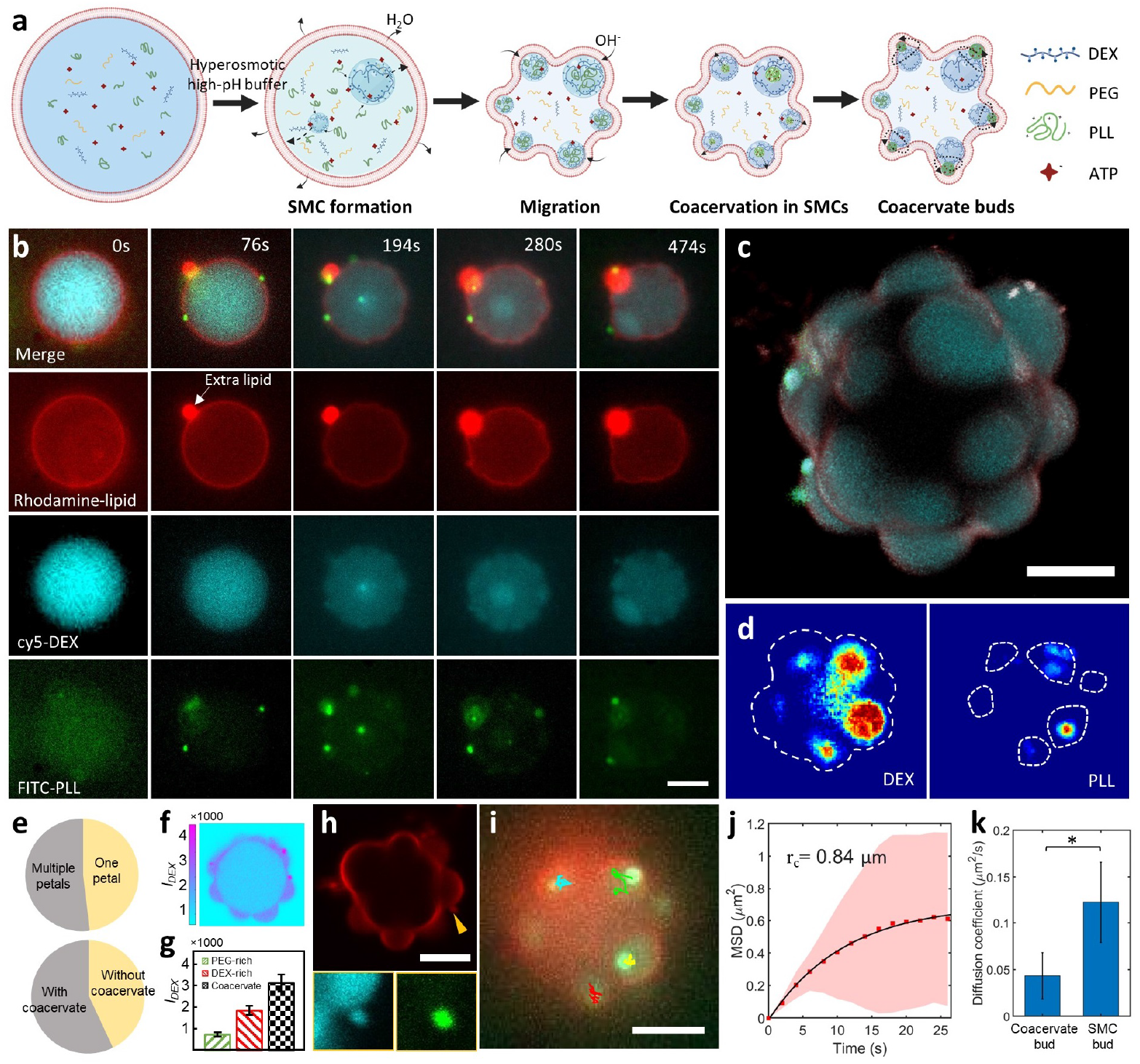
SPS domains regulate coacervates at the membrane. (**a**) Conceptual sketch showing the formation of SPS domains at the membrane, recruitment of PLL, triggering PLL/ATP condensation, and subsequent restricted coacervate motion within the SPS domains. (**b**) Fluorescence time-lapse images showing above-mentioned stages after exposing the liposomes to a hypertonic, high-pH buffer. Volume reduction of the liposome is visible in the lipid channel (in red), sometimes with the extra lipids forming a pocket as indicated. Over time, DEX-rich domains (in cyan) were formed, underwent coalescence, and ultimlately wetted the membrane. These domains recruited PLL molecules and induced coacervation on membrane (observed as intense green dots). (**c**) 3D projection showing the final morphology of a liposome showing petal-shaped SPS domains, some containing condensates. (**d**) Fluorescence-based frequency heat maps showing the observed occurrence frequency (cumulative over one minute) for DEX-rich domains (left) and PLL/ATP condensates (right) in a liposome. The dotted lines are indicative of the lipid membrane (left) and DEX-rich domains (right). (**e**) Pie charts showing the percentage of liposomes with multiple petals (top) and the percentage of petals containing condensates (bottom); *n* = 27 different liposomes. **(f**) Heat map showing DEX distribution in a single plane of the liposome, constructed from a confocal image. (**g**) A bar chart showing dextran-partitioning in PEG-rich phase, DEX-rich phase, and the condensate region; *n* = 5 different liposomes. (**h**) A confocal image of the liposome, showing the budded coacervate region (pointed by the arrow) visualized in lipid channel, while the zoom-ins show the same region in DEX (left) and PLL (right). (**i**) Trajectories of the coacervates showing their restricted motion within the membrane-wetted SMCs. (**j**) MSD plot showing the confined motion of condensates within membrane-bound SMCs; *n* = 4 coacervates within the same liposome. The black curve shows the corraled diffusion fit (R^2^ = 0.99). (**k**) Comparison of the diffusion coefficients between the coacervate buds and the SMC buds (from the previous section); *n* ≥ 4 coacervates; p-value < 0.05. In c-k, the data were collected 20 minutes after the trigger. For all experiments, the lipid composition was 99.9 % DOPC + 0.1% Liss Rhodamine PE (molar ratio). Encapsulating mixture consisted of 7.3 mM DEX (unlabelled: cy5-labeled = 1000:1, molar ratio), 8.6 mM PEG, 2.4 mg/ml PLL (unlabelled: FITC-labeled = 5:1, mass ratio), 0.8 mM ATP, and 15% v/v glycerol in 15 mM citrate-HCl (pH 4). This mixture was combined with an equal volume of feeding aqueous solution containing 500 mM sucrose and 15% v/v glycerol in 15 mM Tris-HCl (pH 9). All scale bar, 5 μm.

To further investigate the regulation of coacervates within a segregative system, we triggered APS and SPS simultaneously by mixing the four components under phase separation conditions, i.e., DEX and PEG concentrations were above the binodal, and the pH (6.3) was suitable for PLL/ATP coacervation. We obtained multi-phase emulsion droplets (Figure 1e), where PLL/ATP droplets were formed and confined within the DEX-rich SMCs, segregated from the PEG-rich environment. We also noted strong partitioning of DEX in the formed condensates, clear from the fluorescence overlap of cy5-labelled DEX and FITC-labelled PLL. We observed the coalescence of SMC domains as well as of the coacervates present inside. As Figure 1f shows, PLL/ATP condensates readily fused with each other upon physical contact, followed by their relaxation into a bigger spherical condensate, indicating their liquid-like behavior within the DEX-rich phase. However, the physical contact was possible only for the coacervates within the same SMC. Figure 1g and Supplementary Movie 1 show the coacervate movement strictly within the SMC domains, even when accentuated by an external fluid flow. Thus, SMC domains acted as incubation chambers for the condensates and physically restricted them.

Next, we used fluorescence recovery after photobleaching (FRAP) to examine the diffusion of coacervate components in such a setting. To check this systematically, we bleached the FITC-PLL fluorescence of entire coacervates in four different environments (Figure 1h): (i) without any DEX or PEG; (ii) in presence of 12 mM DEX; (iii) in presence of 12 mM PEG; and (iv) in presence of 12 mM PEG + 12 mM DEX (SPS conditions). In the first case, we obtained a fluorescence recovery of 85% within 20 minutes (Figure 1h - green panel). The addition of DEX reduced the extent of recovery to almost 50% within the same time period (Figure 1h - red panel). In presence of PEG, the recovery further decreased to only 22% (Figure 1h - black panel). This trend reached its highest extent for the coacervates in SMCs, where only 15% fluorescence was recovered after 20 minutes (Figure 1h - blue panel). The extent of recovery is affected by the PLL concentration in the surrounding dilute phase (*43*), especially since the entire coacervate was bleached. In the case where DEX and/or PEG was present, the decrease in recovery is likely aided by the higher viscosity of the surrounding environment, slowing down the diffusion of PLL molecules. Additionally, the depletion effect by crowding agents has been demonstrated to induce the higher partitioning of PLL in the coacervate phase (*44*), further hindering the recovery. The lowest recovery in the case of coacervates confined within SMCs is a result of concentrated, and thus highly viscous DEX-rich surrounding, and confirms a chemically isolated environment, even for multiple coacervates present within the same SMC. We also observed concomitant differences in the diffusion coefficients of the PLL molecules (*D*_*app,PLL*_). Compared to the control case of no PEG or DEX (1.1 μm2/s), *D*_*app,PLL*_ in PEG and DEX solutions (both ≈ 0.6 μm2/s) was substantially lower (p-value < 0.05, see Figure 1j), likely due to the increased viscosity (*45*). The coacervates confined within the SMC exhibited even lower diffusion coefficient, *D*_*app,PLL*_ ≈ 0.4μm2/s. Thus, these FRAP results indicate that the molecular diffusion between the coacervates and their environment, as well as with neighboring coacervates, is strictly restricted when the coacervates are confined within SMCs.

## APS-SPS multiphase structures within non-membranous microconfinements

Based on the interactions between APS and SPS in bulk, we next turned our attention to whether this hierarchical structure of membraneless compartments would still arise in confined synthetic cell models. We began with applying a simplified synthetic cell model: water-in-oil-in-water double emulsions (Figure 2a). The required APS and SPS components were efficiently encapsulated within double emulsions (osmolarity ≈ 70 mOsm) using a previously established microfluidic platform (*23*) (see Methods for details). The concentration of the SPS components were kept below the binodal (Figure 1b) while the pH was kept at 4.5 in order to prevent coacervation. As Figure 2a-b show, we simultaneously triggered both APS and SPS within the double emulsions using a hypertonic (500 mM sucrose), high pH (8.6) buffer. The hypertonic buffer caused water efflux, decreasing the double emulsion volume, and increasing the concentration of the encapsulated SPS components above the binodal (*22*), while the high pH environment caused proton flux across the boundary, increasing the pH value of the lumen and inducing complex coacervation (*30*). Thus, post-trigger, the homogeneous interior of the double emulsions underwent both APS and SPS, resulting in a multiphase structures. The process was similar to what was observed in bulk settings, in which DEX-rich phases readily fused with each other, eventually forming a single SMC, in which the coacervate was confined. We observed that while some double emulsions first showed SPS and others APS, all of them ultimately formed similar multiphase structure. As seen in bulk experiments (Figure 1e), we observed increased cy5-DEX fluorescence in the coacervate, suggesting a secondary DEX enrichment within the coacervate.

Figure 2c shows the size variation of double emulsions post-trigger. The double emulsions shrunk from ≈ 41 μm to ≈ 38 μm in the first half an hour, and then showed a gradual but still noticeable shrinkage over the next 3 hours before reaching a plateau. Afterward, their size remained constant at ≈ 36.5 μm. Based on the observed volume reduction, concomitant increase in the encapsulated DEX and PEG concentrations were calculated (Figure 2d, see Methods for details). These final concentrations were above the binodal curve, aligning with the observed SMC formation. As shown in Figure 2e, the proportion of double emulsions containing multiphase structures increased rapidly until it plateaued at 70% after 4 hours. We noted that the obtained multiphase structures were not stable over a long-term as the coacervate gradually dissolved over 12 hours (see Supplementary Figure **??**). Nonetheless, we were able to construct a multi-level compartment within double emulsions over the time scale of a several hours, with the MSCs effectively restricting the coacervates.

## SMCs recruit coacervate components to the lipid membrane

To further understand the role of the SPS regulation system in a biomimetic system, we turned to liposomes - aqueous confinements surrounded by a lipid bilayer. We used Octanol-assisted liposome assembly (OLA) (*46, 47*), a liposome generation-visualization microfluidic technique to conduct these experiments (*48*). Based on the well-known interaction between the DEX-rich domain and lipid membranes (*12, 35, 36*), we thought of directing the APS molecules to the membrane via SMCs. As shown in Figure 3a-b, cell-sized (10 − 20 μm in diameter) liposomes were produced, encapsulating the APS and SPS components. To start with, we solely triggered SPS by adding an equal volume of feeding aqueous solution (containing 600 mM sucrose in 25 mM Tris-HCl buffer, pH 9.0) to the liposome suspension (see Materials and Methods). The final pH of the mixture was measured to be 5.1 due to buffering, thus no induction of APS is to be expected. We further demonstrated only SPS formation by triggering the same system in double emulsions, which eliminates any membrane interactions and thus makes it easier to assess the phase separation, and exclusively observing SPS formation (Supplementary Figure **??**). As Figure 3b shows, clear shrinkage in the liposome size was observed after feeding (panels at 384 s), confirming the expected response to the hypertonic environment. Meanwhile, multiple SMCs formed, as evidenced by the increased DEX fluorescence. As expected, there was an overlap of PLL and DEX fluorescence within SMC regions, indicating the recruitment of PLL.

Notably, the SMCs, with PLL recruited inside, readily interacted with the phospholipid membrane. By scanning across different planes, all liposomes within a batch (*n* = 79) were observed to have formed bud-like structures. Figure 3c shows an example, displaying two planes of a liposome 30 minutes post-trigger. In the plane of the SMC bud, marked by a circular lipid structure on the membrane surface, high DEX and PLL fluorescence intensities were observed within the buds. The equatorial plane (insert) shows the non-budded part of the liposome with no SMC localization at the membrane. An intensity profile across the bud (dotted line in Figure 3c), showed a strong peak in both DEX and PLL fluorescence, as well as increased lipid fluorescence (Figure 3d). This overlap of increased lipid fluorescence confirms the pronounced nature of these SMC buds, while being covered by the lipid membrane.

Due to the recruitment within SMCs, PLL molecules were transported to the membrane surface. By plotting the PLL fluorescence intensity as a function of the distance from the center of the liposomes, the PLL fluorescence was indeed observed to be localized at the membrane (Figure 3e, *n* = 10). Furthermore, these PLL-enriched SMCs at the membrane could be seen undergoing fusion events (Figure 3f). As the two SMCs (indicated by white arrows in Figure 3f) came into contact, they merged into a single entity (indicated by green arrows in Figure 3f). These fusion events suggest that the coverage of SMCs by the membrane is incomplete, resulting in protruding buds that are open at the base. Apart from the fusion, the budded SMCs also showed two-dimensional diffusion on the membrane surface (see Supplementary Movie 2). Figure 3g shows the trajectories of several membrane-bound SMCs, restricted along the membrane surface. As seen in Figure 3h, their mean square displacement (MSD) increased linearly in time (R2 = 0.99), indicating an unrestricted two-dimensional (2D) diffusion (*49*). Thus, even if the budded SMCs are coupled to the membrane, they, and as a result the sequestered cargo, are free to diffuse along the membrane.

In conclusion, we demonstrated the transport capability of SMCs for APS components towards the lipid membrane as well as their diffusive motion at the membrane surface. This targeted recruitment is key for triggering coacervation at the membrane, to which we then turned our attention.

## Regulation of coacervates by membrane-interacting SMCs

Based on the SPS-induced PLL migration, we further explored the possibility of inducing coacervation at the membrane (Figure 4a). For this, we applied similar hypertonic conditions by adding equal amounts of feeding aqueous solution (containing 500 mM sucrose in 15 mM Tris-HCl buffer, pH 8.6) to the liposome suspension (see Materials and Methods). The resulting pH after mixing the solutions in equal parts was 6.3, sufficient for APS triggering. This was also demonstrated by triggering both SPS and APS in the double emulsions using the same encapsulated components and triggers (Supplementary Figure **??**). Figure 4b shows fluorescence time-lapse images of the process in liposomes (also see Supplementary Movie 3). As can be seen, both SPS and APS occured within minutes inside the liposome, along with the expected shrinkage of the liposome. The SMCs initially appeared within the liposome interior, after which they wetted the membrane surface and reshaped the liposome into a ‘flower-like’ shape. The SMC domains, representing the ‘petals’, showed strong partitioning of PLL into them. PLL/ATP coacervates were subsequently formed in SMCs and remained close to the membrane. Noticeably, unlike in the case of double emulsions (Figure 2), SMCs with coacervates did not remain suspended in the lumen but ultimately migrated to the membrane surface.

Figure 4c and Supplementary Movie 4 show the 3D rendering of the final morphology of a liposome, captured 20 minutes post-trigger. Multiple DEX-rich petals (in cyan) could be seen, restructuring the liposome into a flower shape. The liposomes showed different morphologies in different planes due to the random distribution of the petals, some of which harbored the formed coacervates (in green) (see Supplementary Movie 5 and Supplementary Figure **??**). Fluorescence-based frequency maps for DEX and PLL further clarified the DEX-rich nature of the petals and the strong confinement experienced by the coacervates within them (Figure 4d, see Methods for details). The formed SMC domains remained quite stable since the attached membrane dynamically arrested their coarsening (*22*). As Figure 4e shows, after 20 minutes of incubation in a high pH, hypertonic environment, 52% of the liposomes (*n* = 27) still maintained multiple ‘petals’ as opposed to a single SMC; 57% of these SMCs contained coacervates.

Similar to the bulk results, DEX fluorescence intensity heat map based on confocal images clearly showed sequential DEX recruitment in the flower-shaped liposomes (Figure 4f). The first recruitment happened in the petals where the DEX intensity was 2.5 times that of the lumen (Figure 4g). Further fluorescence enhancement occurred at the condensates trapped within, showing about 1.7 times further increase in the intensity. As pointed out by the arrow in Figure 4h, the confined coacervates showed a budding behavior, protruding away from the liposome but still surrounded by the lipid membrane. Based on these observations, we concluded that both SMCs and DEX-rich coacervates interact with membranes, forming relatively large petal-like and much smaller bud-like protrusions, respectively.

Importantly, the coacervates were not only restricted by SMCs, and also by their budding behaviour. The individual trajectories in each petal shown in Figure 4i confirm the random but restricted motion of coacervates (see Supplementary Movie 6). Due to the separation of the SMCs, the coacervates remained isolated and could not come in contact with others residing in different petals. We analyzed the diffusive behaviour of the coacervates, based on their MSD. While the MSD trajectory increased linearly at the beginning, it soon plateaued for longer time periods (Figure 4j), indicating this diffusion was free at short times but the effect of barriers became dominant at longer times. Such diffusive motion, exhibiting free movement but within a bounded region, resembled a corralled diffusion (*50*). Indeed, the corresponding equation, (*49*) MSD 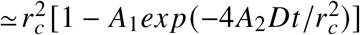, where 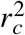 is the corral size, *D* is diffusion coefficient, and *A*_1_ and *A*_2_ are constants determined by the corral geometry, fitted the observed MSD behaviour very well (R2 = 0.99). The corresponding corral size 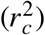 was calculated as 0.7 μ*m*^2^, representing the average restricted area provided by the SPS domain. Comparing the diffusion of these confined coacervates with the membrane-diffusing SMCs (Figure 3) indicated that the diffusion coefficient of the SMC buds was 2.8 times higher than that of the coacervate buds (Figure 4k). This slower diffusion is likely caused by the higher viscosity and the crowded environment in the DEX-rich SMCs.

Thus, we demonstrated APS-membrane interplay with SMCs acting as the mediator between the membrane and the lumen. Such SPS-based regulation of coacervates can prove useful in directed compartmentalization within engineered synthetic cells.

## Discussion and Conclusion

It is well-established that preferential interactions between constituent molecules, either associative or segregative, can destroy homogeneity, drive de-mixing, and produce co-existing phases within cellular systems. But how these two types of interactions, occurring simultaneously and dynamically within the cytoplasmic space, influence each other is still largely unknown. Using a model system consisting of four components in synthetic giant vesicles, we have demonstrated that SPS can regulate APS at both molecular and coacervate level. We further showcase that the APS-SPS-membrane interplay can generate an entirely new degree of freedom regulating dynamic reorganization, spatial localization, movement restriction, and surface interactions of condensates within the crowded aqueous space.

We show molecular enrichment via SPS-based membraneless confinements (SMCs) recruiting and transporting molecules to the membrane (or keeping them in the lumen), allowing further condensation under the right conditions. Such membrane-targeted migration of molecules could improve the efficiency of on-membrane or transmembrane reactions and it will be beneficial to assess the generality of such affinity-based compartmentalization in diverse chemical systems including elastin-like (*51, 52*) and other polypeptides (*41, 53*), RNA (*54*), or even nanoliposomes (*55*). Interestingly, SMCs have also been demonstrated to partition non-biorelevant components like metal particles (*56*) or organic polymers (*57*). These substances, which normally do not exist in living organisms, may bring new functionalities to synthetic cells (*58*).

We further demonstrated that SMCs can spatially restrict the coacervate movement. Moreover, due to the lack of molecular communication across coacervates (*25*), even within the same SMC, each coacervate can be considered independent. Therefore, for reaction cascades, these coacervate-in-SMC multiphases may be a good strategy for providing a membrane-free domain. Also, as the observed SPS is mainly driven by concentration and is stable across a wider pH range, one can easily switch the electrostatic environment to tune APS individually (*59*). Such hierarchical multiphase assembly with distinct triggers allows for a more flexible and dynamic control.

While high-throughput single emulsion microfluidics (*60*) provides an efficient solution for studying sub-compartmentalization (*61*), the phase separation largely depends on the encapsulated components rather than the subsequent external stimuli (*62*), making the single emulsion system relatively inflexible. The double emulsion system we used here provides the ability to trigger processes within the containers simply by tuning the extracellular environment (*23*). Notably, double emulsions are robust in various physicochemical variations including pH, temperature and osmotic pressure (*23*), making them a handy system if membrane interactions are not necessary or to be deliberately avoided.

When it comes to the use of liposomes, APS-membrane interactions are facilitated via SMCs. With existing coacervate-membrane binding usually relying on conjugations including charge-based (*29*) and hydrophobic (*30*) interactions, SMCs-based delivery shown here can be considered a conjugation-free strategy. Furthermore, more often than not, APS/SPS interactions with the membrane are studied from the outer leaflet, partly owing to the difficulty of encapsulation (*29, 33, 34*). Because of our microfluidic approach, we are able to study them from inside the liposomes, allowing us to mimic a more natural system.

In synthetic cell confinements, the isolation of coacervates as individual reaction hubs has high significance but still requires exploration. Here, the membrane-bound SMCs offer microchambers for the co-existence of multiple similar condensates within a single liposome. As SMCs can be stable for hours without fusion, they can provide independent compartments for their subordinate coacervates, and this stability could be further prolonged using lipids with different fluidities (*22*). Taken together, our results highlight a heretofore unappreciated factor producing new functional levels of organization in cells. Given the vast molecular diversity of the cytoplasmic crowd, which inevitably results in both associative and segregative tendencies, our findings suggest a universal physicochemical principle and a non-specific biological strategy to spatially and temporally regulate biomolecular condensates. This approach has broad applications in controlling the behavior of membraneless organelles and construct diverse synthetic cell architectures.

## Acknowledgments

C.C. acknowledges financial support from the graduate school (VLAG) to support scientific travelling. Schematics were created using BioRender.com.

## Author contributions

Conceptualization: C.C., C. L., C. F. C., A. N. P., and S.D.; Investigations and methodology: C.C., C. L., C. F. C., A. N. P., and S.D.; bulk experiments – C.C., C. L., C. F. C.; microfluidics and microscopy – C.C., K.A.G, Data Curation: C.C., C. L., C. F. C.; Formal Analysis: C.C., C. L., C. F. C., S.D.; Funding Acquisition and project administration: A.N.P., S.D.; Resources: A. N. P, S.D.; Supervision and validation: A. N. P., S.D.; Visualization: C.C., S.D.; Writing (original draft): C.C, S.D.; Writing (review and editing): C.C., C. L., C. F. C., K.A.G, A. N. P., and S.D. All authors have read and given their consent to the final version of the publication.

## Data and materials availability

Data supporting the findings of this study are available within the paper, its Supplementary Information, Supplementary Movie 1–6, and Source Data 1–2. Any additional supporting data are available from the corresponding authors upon reasonable request. The MATLAB codes used for image processing (Fig. 3e, g, h and Fig. 4d, f, i, j) are provided as Source Data 2.

## Statistics and reproducibility

Experiments related to the binodal curve (Fig. 1b, Supplementary Fig. S1) were repeated once for each of the five conditions. Experiments related to PLL/ATP partitioning (Fig. 1c-d), APS-SPS dynamics (Fig. 1e-g, with varied ATP concentrations), FRAP experiments (Fig. 1h-j), double emulsions (Fig. 2, Supplementary Fig. S2 and S4), and liposomes (Fig. 3, with varied osmotic pressure strengths between 300-600 mOsm; Fig. 4) were repeated at least three times with similar results. Experiments related to SPS triggering in double emulsions (Supplementary Fig. S3) were repeated once.

## Supplementary materials

Materials and Methods

Figs. S1 to S5

Tables S1

Movie S1 to S6

Data S1 to S2

## References

1. C. Xu, N. Martin, M. Li, S. Mann, Living material assembly of bacteriogenic protocells. Nature 609 (7929), 1029–1037 (2022).

2. D. Bracha, M. T. Walls, C. P. Brangwynne, Probing and engineering liquid-phase organelles. Nature biotechnology 37 (12), 1435–1445 (2019).

3. W. Mu, et al., Superstructural ordering in self-sorting coacervate-based protocell networks. Nature Chemistry 16 (2), 158–167 (2024).

4. S. C. Shetty, et al., Directed signaling cascades in monodisperse artificial eukaryotic cells. ACS nano 15 (10), 15656–15666 (2021).

5. S. Deshpande, W. K. Spoelstra, M. Van Doorn, J. Kerssemakers, C. Dekker, Mechanical division of cell-sized liposomes. ACS nano 12 (3), 2560–2568 (2018).

6. S. Deshpande, S. Wunnava, D. Hueting, C. Dekker, Membrane tension–mediated growth of liposomes. Small 15 (38), 1902898 (2019).

7. R. Dimova, C. Marques, The giant vesicle book (2019).

8. N.-N. Deng, W. T. Huck, Microfluidic formation of monodisperse coacervate organelles in liposomes. Angewandte Chemie 129 (33), 9868–9872 (2017).

9. S. Deshpande, et al., Spatiotemporal control of coacervate formation within liposomes. Nature communications 10 (1), 1800 (2019).

10. Z. Lin, T. Beneyton, J.-C. Baret, N. Martin, Coacervate Droplets for Synthetic Cells. Small Methods 7 (12), 2300496 (2023).

11. K. A. Ganar, L. Leijten, S. Deshpande, Actinosomes: condensate-templated containers for engineering synthetic cells. ACS synthetic biology 11 (8), 2869–2879 (2022).

12. L. B. Case, X. Zhang, J. A. Ditlev, M. K. Rosen, Stoichiometry controls activity of phase-separated clusters of actin signaling proteins. Science 363 (6431), 1093–1097 (2019).

13. S. An, R. Kumar, E. D. Sheets, S. J. Benkovic, Reversible compartmentalization of de novo purine biosynthetic complexes in living cells. Science 320 (5872), 103–106 (2008).

14. J. B. Woodruff, et al., The centrosome is a selective condensate that nucleates microtubules by concentrating tubulin. Cell 169 (6), 1066–1077 (2017).

15. E. Boke, et al., Amyloid-like self-assembly of a cellular compartment. Cell 166 (3), 637–650 (2016).

16. S. Deshpande, C. Dekker, Studying phase separation in confinement. Current Opinion in Colloid & Interface Science 52, 101419 (2021).

17. C. P. Brangwynne, et al., Germline P granules are liquid droplets that localize by controlled dissolution/condensation. Science 324 (5935), 1729–1732 (2009).

18. M. Feric, et al., Coexisting liquid phases underlie nucleolar subcompartments. Cell 165 (7), 1686–1697 (2016).

19. S. Zhu, et al., Demixing is a default process for biological condensates formed via phase separation. Science 384 (6698), 920–928 (2024).

20. A. W. Folkmann, A. Putnam, C. F. Lee, G. Seydoux, Regulation of biomolecular condensates by interfacial protein clusters. Science 373 (6560), 1218–1224 (2021).

21. M. van Dop, et al., DIX domain polymerization drives assembly of plant cell polarity complexes. Cell 180 (3), 427–439 (2020).

22. W.-C. Su, et al., Kinetic control of shape deformations and membrane phase separation inside giant vesicles. Nature Chemistry 16 (1), 54–62 (2024).

23. C. Chen, et al., Elastin-like polypeptide coacervates as reversibly triggerable compartments for synthetic cells. Communications Chemistry 7 (1), 1–11 (2024).

24. M. P. Tran, et al., A DNA segregation module for synthetic cells. Small 19 (13), 2202711 (2023).

25. Q.-H. Zhao, F.-H. Cao, Z.-H. Luo, W. T. Huck, N.-N. Deng, Photoswitchable Molecular Communication between Programmable DNA-Based Artificial Membraneless Organelles. Angewandte Chemie 134 (14), e202117500 (2022).

26. T. Beneyton, C. Love, M. Girault, T.-Y. D. Tang, J.-C. Baret, High-throughput synthesis and screening of functional coacervates using microfluidics. ChemSystemsChem 2 (6), e2000022 (2020).

27. K. A. Ganar, L. W. Honaker, S. Deshpande, Shaping synthetic cells through cytoskeleton-condensate-membrane interactions. Current Opinion in Colloid & Interface Science 54, 101459 (2021).

28. B. Wang, et al., Liquid–liquid phase separation in human health and diseases. Signal Transduction and Targeted Therapy 6 (1), 290 (2021).

29. T. Lu, et al., Endocytosis of coacervates into liposomes. Journal of the American Chemical Society 144 (30), 13451–13455 (2022).

30. M. G. Last, S. Deshpande, C. Dekker, pH-controlled coacervate–membrane interactions within liposomes. ACS nano 14 (4), 4487–4498 (2020).

31. C. Bussi, et al., Stress granules plug and stabilize damaged endolysosomal membranes. Nature 623 (7989), 1062–1069 (2023).

32. A. Mangiarotti, R. Dimova, Biomolecular Condensates in Contact with Membranes. Annual Review of Biophysics 53 (2024).

33. A. Mangiarotti, et al., Biomolecular condensates modulate membrane lipid packing and hydration. Nature Communications 14 (1), 6081 (2023).

34. T. Lu, X. Hu, M. H. van Haren, E. Spruijt, W. T. Huck, Structure-Property Relationships Governing Membrane-Penetrating Behaviour of Complex Coacervates. Small p. 2303138 (2023).

35. R. Dimova, R. Lipowsky, Giant Vesicles Exposed to Aqueous Two-Phase Systems: Membrane Wetting, Budding Processes, and Spontaneous Tubulation. Advanced Materials Interfaces 4 (1), 1600451 (2017).

36. M. R. Helfrich, L. K. Mangeney-Slavin, M. S. Long, K. Y. Djoko, C. D. Keating, Aqueous phase separation in giant vesicles. Journal of the American Chemical Society 124 (45), 13374–13375 (2002).

37. Y. Li, R. Lipowsky, R. Dimova, Membrane nanotubes induced by aqueous phase separation and stabilized by spontaneous curvature. Proceedings of the National Academy of Sciences 108 (12), 4731–4736 (2011).

38. Y. Li, R. Lipowsky, R. Dimova, Transition from complete to partial wetting within membrane compartments. Journal of the American Chemical Society 130 (37), 12252–12253 (2008).

39. Y. Liu, J. Agudo-Canalejo, A. Grafmuller, R. Dimova, R. Lipowsky, Patterns of flexible nanotubes formed by liquid-ordered and liquid-disordered membranes. Acs Nano 10 (1), 463–474 (2016).

40. M. Andes-Koback, C. D. Keating, Complete budding and asymmetric division of primitive model cells to produce daughter vesicles with different interior and membrane compositions. Journal of the American Chemical Society 133 (24), 9545–9555 (2011).

41. M. S. Long, A.-S. Cans, C. D. Keating, Budding and asymmetric protein microcompartmentation in giant vesicles containing two aqueous phases. Journal of the American Chemical Society 130 (2), 756–762 (2008).

42. Y. Li, H. Kusumaatmaja, R. Lipowsky, R. Dimova, Wetting-induced budding of vesicles in contact with several aqueous phases. The Journal of Physical Chemistry B 116 (6), 1819–1823 (2012).

43. W. M. Aumiller Jr, F. Pir Cakmak, B. W. Davis, C. D. Keating, RNA-based coacervates as a model for membraneless organelles: formation, properties, and interfacial liposome assembly. Langmuir 32 (39), 10042–10053 (2016).

44. S. Park, et al., Dehydration entropy drives liquid-liquid phase separation by molecular crowding. Communications Chemistry 3 (1), 83 (2020).

45. I. Avramov, Relationship between diffusion, self-diffusion and viscosity. Journal of Non-Crystalline Solids 355 (10-12), 745–747 (2009).

46. S. Deshpande, Y. Caspi, A. E. Meijering, C. Dekker, Octanol-assisted liposome assembly on chip. Nature communications 7 (1), 10447 (2016).

47. S. Deshpande, C. Dekker, On-chip microfluidic production of cell-sized liposomes. Nature protocols 13 (5), 856–874 (2018).

48. C. Chen, K. A. Ganar, S. Deshpande, On-Chip Octanol-Assisted Liposome Assembly for Bioengineering. JoVE (Journal of Visualized Experiments) (193), e65032 (2023).

49. M. J. Saxton, K. Jacobson, Single-particle tracking: applications to membrane dynamics. Annual review of biophysics and biomolecular structure 26 (1), 373–399 (1997).

50. Y. Chen, B. C. Lagerholm, B. Yang, K. Jacobson, Methods to measure the lateral diffusion of membrane lipids and proteins. Methods 39 (2), 147–153 (2006).

51. H. Zhao, et al., Spatiotemporal Dynamic Assembly/Disassembly of Organelle-Mimics Based on Intrinsically Disordered Protein-Polymer Conjugates. Advanced Science 8 (24), 2102508 (2021).

52. H. Zhao, V. Ibrahimova, E. Garanger, S. Lecommandoux, Dynamic spatial formation and distribution of intrinsically disordered protein droplets in macromolecularly crowded protocells. Angewandte Chemie International Edition 59 (27), 11028–11036 (2020).

53. M. S. Long, C. D. Jones, M. R. Helfrich, L. K. Mangeney-Slavin, C. D. Keating, Dynamic microcompartmentation in synthetic cells. Proceedings of the National Academy of Sciences 102 (17), 5920–5925 (2005).

54. C. Qi, et al., Multicompartmental coacervate-based protocell by spontaneous droplet evaporation. Nature Communications 15 (1), 1107 (2024).

55. H. Seo, C. Nam, E. Kim, J. Son, H. Lee, Aqueous two-phase system (ATPS)-based polymersomes for particle isolation and separation. ACS Applied Materials & Interfaces 12 (49), 55467–55475 (2020).

56. M. R. Helfrich, M. El-Kouedi, M. R. Etherton, C. D. Keating, Partitioning and assembly of metal particles and their bioconjugates in aqueous two-phase systems. Langmuir 21 (18), 8478–8486 (2005).

57. L. W. Honaker, et al., 2D and 3D Self-Assembly of Fluorine-Free Pillar-[5]-Arenes and Perfluorinated Diacids at All-Aqueous Interfaces. Advanced Science p. 2401807 (2024).

58. H. R. Vutukuri, et al., Active particles induce large shape deformations in giant lipid vesicles. Nature 586 (7827), 52–56 (2020).

59. J. Li, et al., Programmable spatial organization of liquid-phase condensations. Chem 8 (3), 784–800 (2022).

60. N. Appelman, C. Chen, I. Gruppen, S. Deshpande, Interface-Driven Spontaneous Differentiation-Repulsion Behavior in Isochemical Droplet Populations. Advanced Materials Interfaces 11 (9), 2300617 (2024).

61. Y. Cao, et al., Partitioning-induced isolation of analyte and analysis via multiscaled aqueous two-phase system. Analytical Chemistry 95 (10), 4644–4652 (2023).

62. F. Chen, et al., Size Scaling of Condensates in Multicomponent Phase Separation. Journal of the American Chemical Society (2024).

