## Supplementary Information for "Regulating biocondensates within synthetic cells via segregative phase separation"

**This PDF file includes:**

Materials and Methods

Figures S1 to S5

Tables S1

Captions for Movies S1 to S6

Captions for Data S1 to S2

**Other Supplementary Materials for this manuscript:**

Movies S1 to S6

Data S1 to S2

### Materials and Methods

#### Materials

Dextran (MW 9-11 kDa, No. D9260), Polyethylene glycol (MW 8 kDa, No. 89510), Poly-L-lysine (MW 15-30 kDa, P7890), Poly-L-lysine–FITC labeled (MW 15-30 kDa, P3543), Adenosine 5'-triphosphate disodium salt hydrate (ATP, A2383), sucrose (S0389), Tris(hydroxymethyl)aminomethane (Tris-base, 252859), sodium citrate tribasic dihydrate (citrate-base, 71405), polyvinyl alcohol (PVA, average MW 30-70 kDa, P8136), 1-Octanol (297887), glycerol (G2025), Pluronic™ F-68 non-ionic surfactant (24040032) and hydrochloric acid were purchased from Sigma-Aldrich. Labeled dextran (Alexa Fluor™ 647, 10,000 MW, D22914) was purchased from Fischer Scientific B.V. Phospholipids including 18:1 (Δ9-Cis) PC (DOPC) (SKU 850375C) and 18:1 Liss Rhod PE (SKU 810150C) were purchased from Avanti Polar Lipids, Inc. $N^6$-(6-Aminohexyl)-adenosine-5'-triphosphate labeled with cy3 (NU-805-CY3) was purchased from Jena Bioscience. Sylgard™ 184 silicone elastomer (PDMS) and curing agent were purchased from Dow. Silicon wafer was bought from Silicon Materials. Photoresist (EpoCore 10) and photoresist developer (mr-Dev 600) were purchased from Micro resist technology GmbH. Microfluidic accessories including liquid flows tygon tubing coil 1/16" OD X 0.02" ID (SKU: LVF-KTU-13), stainless steel 90° Bent PDMS couplers (SKU: PN-BEN-23G), rapid-core microfluidic punches (D= 0.5mm and 3mm), tygon® tubing 1/16" ODx 0.02" and HFE 7500 fluorinated oil containing 2% FluoSurf-C surfactant were purchased from Darwin Microfluidics. Elveflow pressure controller OB1-MK3 was used to control the fluid flow.

#### Stock solutions

Stock solutions of DEX (30 mM), PEG (70 mM) and sucrose (1 M) were prepared by dissolving the respective materials in Milli-Q water using a volumetric flask. On average, 40 mM of DEX 10k is equivalent to ~34 % w/v while 70 mM of PEG 8k is equivalent to ~51 % w/v.

The lipid stock solutions were prepared as described in detail previously (*?*). In this paper, we used a mixture of DOPC and of Liss Rhod PE (molar ratio = 1000:1) and the stock concentration for DOPC was 100mg/ml. Briefly, we pipetted appropriate volume of DOPC and Liss Rhod PE chloroform solutions at the bottom of the round-bottom flask. The chloroform was fully evaporated by passing a gentle stream of nitrogen into the flask and desiccating the flask for more than two hours to form a dry lipid film at the bottom. The 10% (w/v) lipid stock was then made by dissolving the lipids in appropriate volume of ethanol. For long-term preservation, the stock was stored in a dark glass vial filled with an inert atmosphere at -20°C.

#### Elucidation of the phase diagram

The binodals of PEG and DEX were measured at room temperature using a variation of cloud-point titration (**?**). A known volume of DEX stock solution was added to a small glass vial. A small known volume of PEG stock solution was then added and the mixture was stirred. This process was repeated until the resulting mixture reached its cloud-point and became turbid. Milli-Q water was then added in small volume increments to the mixture, followed by mixing, until the mixture became clear again. Alternating volumes of the PEG stock and milli-Q water were titrated into the mixture, crossing above and below the cloud-point, until the cloud point could not be reached or a significantly large volume of the PEG was required. The process was then repeated switching DEX and PEG. Subsequent binodals that include different buffers or glycerol were produced by adding equal concentrations to each of the stock solutions and water so the concentration of the added component would remain constant. The procedure was otherwise identical.

#### Measurement of partition coefficients

The partitioning of PLL or ATP in a phase-separated DEX/PEG mixture was measured by mixing corresponding components in Eppendorf tubes. Each mixture consisted of PEG and DEX in varying concentrations in 25 mM Tris-HCl at pH 7.4 and with either 2.4 mg/mL PLL (unlabelled: FITC-labeled = 10:1, mass ratio) or 0.8 mM ATP (unlabelled: cy3-labeled = 1000:1, molar ratio). The concentration of PEG was varied between 20, 15, and 10 wt% while DEX varied between 15, 10, and 5 wt%, resulting in nine total combinations of PEG and DEX. These vials were vortexed to ensure mixing and then left in the fridge overnight to allow the PEG and DEX to phase separate. On a clean glass slide, 5 uL droplets were placed from either the top PEG-rich phase or the bottom DEX-rich phase and immediately imaged using fluorescence microscopy. The average fluorescence intensity was normalized as following (**?**):

$$I = I_{measured} - I_{reference},$$

where $I$ is the normalized intensity, $I_{measured}$ is the measured intensity in droplet and $I_{reference}$ is the mean intensity in a random area outside the droplet. The partitioning coefficients of the fluorescent components, $K_{PLL}$ or $K_{ATP}$, were calculated by:

$$K = \frac{I_{DEX}}{I_{PEG}},$$

where $I_{DEX}$ and $I_{PEG}$ is the normalized intensity of PLL/ATP in the DEX-rich phase and the PEG-rich phase, respectively. Each data point had three replicates.

**Microfabrication and surface functionalization**

Master wafers were prepared according to the previously described microfabrication method (*?*) and the channel height was kept at 10 $\mu$m. Microfluidic devices were prepared by standard soft lithography method (*?, ?, ?*). Briefly, PDMS and the curing agent were mixed in a 10:1 weight ratio and then poured on the master wafer and degassed using a vacuum desiccator. Meanwhile, a PDMS-coated glass coverslip (Corning® no. 1) was coated with PDMS by spin coating at 500 rpm for 15 s (at an increment of 100 rpm/s) and then at 1000 rpm for 30 s (at an increment of 500 rpm/s). Both PDMS-covered wafer and PDMS-coated glass were baked at 70 °C for 2 hours. The hardened PDMS block was carefully removed, and inlets and outlet holes were punched using a biopsy punch of diameter 0.3 mm. The PDMS block was then bonded on the coverslip after 30 seconds of plasma treatment at 12 MHz (RF mode high) using a plasma cleaner (Harrick Plasma PDC-32G). The bonded device was then baked at 70 °C for two hours.

After baking, the production chip underwent a PVA (5% w/v, molecular weight 30 -70 kDa) treatment as described previously (*?*). Briefly, the outer aqueous inlet was flowed with a PVA solution whereas the inner aqueous and oil inlets were kept at a positive pressure to retain the PVA-air boundary stable at the second production junction. After 15 minutes of incubation, PVA solution was pushed out by applying maximum pressure (2 bar) on the inner aqueous and oil inlets and the extra liquid was removed by applying a negative pressure (-1 bar) at the exit. The device was then dried for 15 minutes by baking on a hot plate at 120 °C.

PDMS wells for FRAP and double emulsion experiments were prepared using a clean silicon wafer without any pattern. First, non-patterned PDMS blocks were prepared similarly to the description above by pouring PDMS-curing mixture over the wafer, followed by baking and removing the cured material from the wafer. The blocks were then punched for 5 mm-holes as the wells. After plasma bonding on a PDMS-coated glass slide, the wells were surface-functionalized by pipetting PVA solution follwing by 15-minute incubation. The PVA was then pipetted out and the device was baked on a hot plate at 120 °C for 15 minutes. After surface functionalization, all devices were stored at room temperature and can be stable for months.

**Bulk experiments**

In the bulk SPS-APS interaction experiment (Figure 1 f-g), we made a mixture of 3 mg/ml PLL (unlabelled: FITC-labeled = 5:1, mass ratio), 4 mM ATP, 12 mM DEX, 12 mM PEG and 15% glycerol in a buffer made by one-to-one mixing of 15 mM Tris-HCl (pH 9) and 15 mM citrate-HCl (pH 4) in Eppendorf tube. After proper mixing by pipetting, 5 $\mu$L solution was dropped on a cover glass and observed under the fluorescence microscope. A transparent lid was used to prevent evaporation.

For FRAP experiments (Figure 1 h-j), we made four mixtures containing 3 mg/ml PLL (unlabelled: FITC-labeled = 5:1, mass ratio), 4 mM ATP and 15% glycerol in a buffer made by one-to-one mixing of 15 mM Tris-HCl (pH 9) and 15 mM citrate-HCl (pH 4) with in Eppendorf tubes: 1) no crowding agent, 2) 12 mM DEX, 3) 12 mM PEG, 4) 12 mM DEX and 12 mM PEG. After proper mixing by pipetting, 10 $\mu$L solution was dropped on a PDMS well with cover glass and observed under the fluorescence microscope. For bleaching, the region of interest (ROI) were entire condensates of approximately 14 $\mu$m in diameter. Intensity of the bleached area was normalized using the equation:

$$f(t) = \frac{I_{\text{correct}}(t) - \min(I_{\text{correct}})}{I_{\text{correct}}(0) - \min(I_{\text{correct}})},$$

where

$$I_{correct} = C(t) * I(t),$$

and

$$C(t) = \frac{R(0)}{R(t)}.$$

*R(t)* and *I(t)* indicate the fluorescence intensity of the reference droplet at time *t* and the original fluorescence intensity of the bleached region at time *t*, respectively; $\min(I_{correct})$ indicates the minimum value of $I_{correct}$, which is obtained right after the sample is bleached (***?***). The normalized intensity was fitted using the function:

$$f(t) = A(1 - e^{(-t/\tau)}),$$

where $A$ and $\tau$ indicate the amplitude of recovery and the relaxation time, respectively. The apparent diffusion coefficient ($D_{app}$) was calculated using the formula:

$$D_{app} \simeq \frac{\omega^2}{t_{(\frac{1}{2})}},$$

where $t_{(1/2)}$ is the half-life fluorescence recovery and $\omega^2$ is the area of the bleached cross section. The half-life $t_{(1/2)}$ was calculated using the formula:

$$t_{(1/2)} = \ln(2)\tau.$$

**Liposome and double emulsion production**

Octanol-assisted liposome assembly (OLA) method (**?**) was applied for the liposome production. Four solutions were prepared: inner aqueous (IA), outer aqueous (OA), lipids in 1-octanol (LO), and exit well aqueous (EA). IA, OA, and EA always contained 15% v/v glycerol and a pH-regulating critrate-HCl buffer (pH ≈ 4). Additionally, 5% w/v F68 surfactant was always present in OA. Phase separation components coexisted in IA as a homogeneous solution: 7.3 mM DEX (unlabelled: cy5-labeled = 1000:1, molar ratio), 8.6 mM PEG, 2.4 mg/ml PLL (unlabelled: FITC-labeled = 5:1, mass ratio), and 0.8 mM ATP. The osmolarity of the aqueous solutions was balanced by the addition of 70 mM sucrose in OA and EA. The lipid-carrying organic phase was prepared by mixing 10% lipid stock (10% DOPC and 0.1% Liss Rhod PE, w/v in ethanol) with 1-octanol to a final concentration of 0.2% w/v. A detailed protocal is described elsewhere (*?*). After adjusting the three inlet pressures and obtaining stable production, the open well was filled with 10 $\mu$L EA to collect the liposomes. The production phase normally lasted for about 30 minutes.

Double emulsion production was conducted by a double-junction microfluidic design as described previously (**?**). We used the same IA and OA components as the liposome experiments. The oil phase was prepared by mixing labelled-lipid stock (0.1% Liss Rhod PE, w/v in ethanol) with HFE 7500 fluorinated oil (containing 2% FluoSurf-C surfactant) to a final concentration of 0.002% w/v. A clean pipette tip (200 $\mu$L) was inserted into the outlet to collect the produced double emulsions. The dispersion was stored in amber-colored bottles and kept at 4 °C.

### On-chip treatment and observation

In the liposome experiments, triggering was conducted in the same chip as the production chip. Briefly, both APS and SPS were triggered simultaneously by gently removing 5 $\mu$L of the solution so as to not disturb the settled liposomes and refilling the well with 5 $\mu$L of feeding aqueous (FA). The well was covered with a coverslip to prevent evaporation, except during solution exchange.

In the double emulsion experiments, the triggering was performed by replacing the external solution in which the double emulsions were dispersed. Initially, the PDMS well was filled by pipetting 90 $\mu$L of OA and 10 $\mu$L of double emulsion suspension. After the double emulsions sank to the bottom, 50 $\mu$L of the supernatant was removed, followed by immediately refilling with 50 $\mu$L of FA. The well was consistently covered with a coverslip except during solution exchange.

A detailed list of IA, OA, EA and FA solution compositions for various experiments can be found in Supplementary Table 1.

### Binodal crossing analysis

The binodal curve was fitted based on all experimental data in varied conditions (pH, viscosity, and ionic strengths) using the equation,

$$c_{\mathrm{DEX}} = a + b * c_{\mathrm{PEG}} + d * c_{\mathrm{PEG}}^2$$

where $a, b, c$ are the fitting constants, $c_{DEX}$ and $c_{PEG}$ are the concentrations of DEX and PEG respectively. Based on the fitted binodal curve, we further calculated the DEX and PEG concentrations after triggering. First, the volume ratio $R$ was obtained as

$$R = (\frac{D_{\mathrm{before}}}{D_{\mathrm{after}}})^3$$

where $D_{before}$ and $D_{after}$ are the average diameters of the liposomes before hypertonic high-pH feeding and 4 hours post-feeding, respectively. The DEX and PEG concentrations inside the liposomes were then calculated as

$$c_{\mathrm{after}} = c_{\mathrm{before}} * R$$

where $c_{before}$ and $c_{after}$ are the concentrations of DEX and PEG before and after hypertonic high-pH trigger respectively.

### Microscopy

Images for bulk and on-chip experiments were acquired using a Nikon-Ti2-Eclipse inverted fluorescence microscope equipped with pE-300ultra illumination system, Nikon Plan F 10x (numerical aperture, NA 0.3) objective, Nikon Plan Fluor 40x (NA 1.30) oil objective or Nikon Plan Apo 100x (NA 1.45) oil objective, and appropriate filter sets (Semrock). The excitation wavelength are 460 nm for FITC and 550 nm for Rhodamine and cy5. In case of fluorescence visualization, samples were excited using 2-10% light intensity and an exposure time of 10-100 ms. All images were acquired using a Prime BSI Express sCMOS camera.

FRAP experiments were performed on Leica SP8-SMD microscope using 63x (NA 1.2) water objective. For bleaching, the ROI was bleached using 100% laser intensity for 5.1 seconds and recovery of the bleached area was recorded for every 0.5 seconds for approximately 20 minutes.

Confocal images were acquired using a Nikon C2 Confocal laser scanning microscope equipped with Ti2 Illuminator-DIA system, Nikon Plan Apo 60x (NA 1.4) oil objective, and appropriate filter sets (Semrock).

### Image analysis

ImageJ was used for image processing and analysis in the case of FRAP experiments, fluorescence intensity analysis as well as size measurements. In case of double emulsion experiments, only those double emulsions that were non-clustered and in focus were taken into account for the analysis (MATLAB R2019b) Error bars in the graphs indicate the standard deviation of the mean for respective samples.

For PLL distribution along the radius, the membrane boundary was selected based on the lipid channel, and the center of the liposome was calculated. Subsequently, PLL fluorescence intensity from the center to the boundary was measured, normalized by the maximum intensity value, and plotted against normalized liposome radius. In case of DEX/PLL fluorescence appearance frequency heat map (Figure 4d), pixels with an intensity higher than 150 were considered as an appearance of MSCs/coacervates in that frame. All frames in the video (1 min duration) were analyzed and the cumulative number of occurrences in each pixel was summarized and expressed as the frequency heat map. For DEX fluorescence distribution in one liposome (Figure 4f), the fluorescence intensity in each point for a single time frame was summarized and expressed by a heat map.

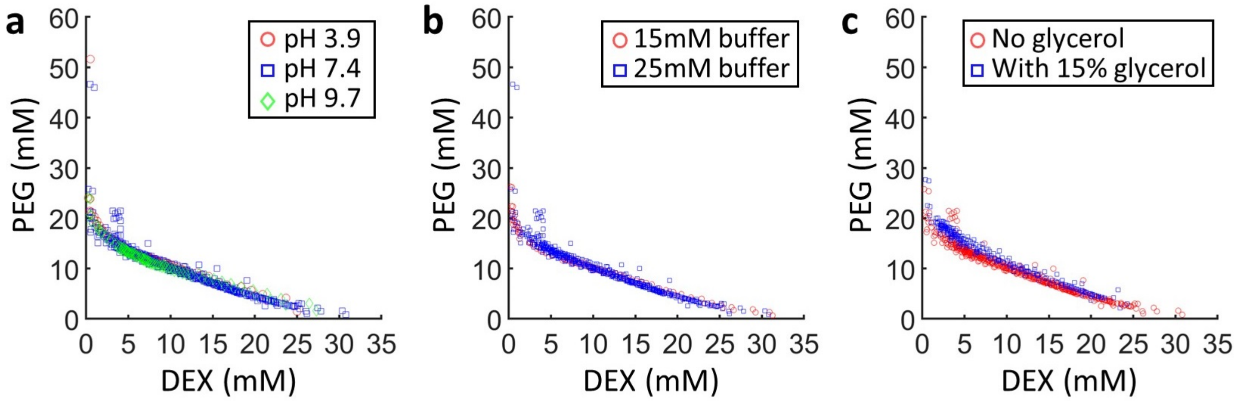


**Figure S1**: **Binodal curves for PEG (8 kDa) and DEX (10 kDa) at room temperature for three different parameter variations.** The binodal line is obtained by the cloud-point method in bulk solution in varied (a) pH, (b) buffer concentrations, and (c) viscosity.

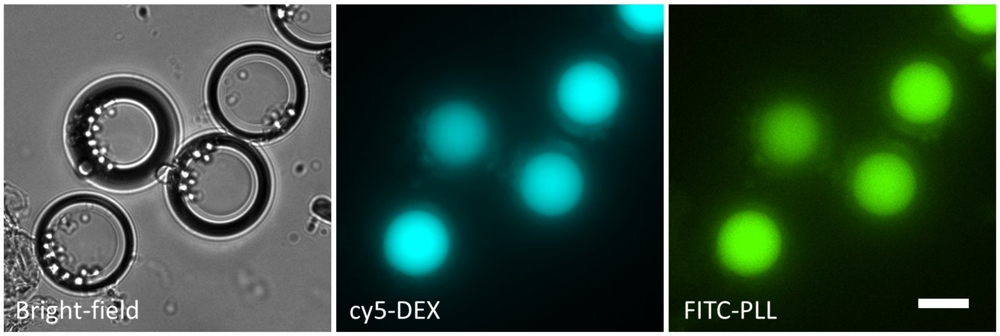


**Figure S2**: **Coacervates confined within SMCs, formed within double emulsions, tend to dissolve over longer duration.** Bright-field and corresponding fluorescence images, captured 13 hours after the hypertonic, high-pH trigger. The bright-field image shows a single SMC domain in each double emulsion, as confirmed by the DEX fluorescence. The PLL fluorescence shows complete overlap with DEX fluorescence, but no higher-fluorescence regions are present witihn the SMC domain, indicating coacervation dissolution. Encapsulating mixture consisted of 7.3 mM DEX (unlabelled: cy5-labeled = 1000:1, molar ratio), 8.6 mM PEG, 2.4 mg/ml PLL (unlabelled: FITC-labeled = 5:1, mass ratio), 0.8 mM ATP, and 15% v/v glycerol in 15 mM citrate-HCl (pH 4). This mixture was combined with an equal volume of feeding aqueous solution containing 500 mM sucrose and 15% v/v glycerol in 15 mM Tris-HCl (pH 9). Scale bar, 20 µm.

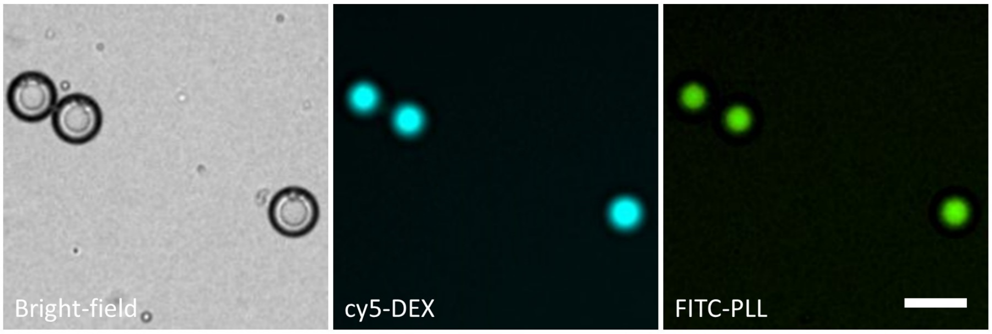


**Figure S3**: **Exposing double emulsions to a hypertonic, high-pH buffer with a higher ionic strength (25 mM Tris-HCl) does not lead to coacervation but only SMC formation.** Bright-field and corresponding fluorescence images, captured 2 hours after the trigger. The bright-field image shows a single SMC domain in each double emulsion, as confirmed by the DEX fluorescence. The PLL fluorescence shows complete overlap with DEX fluorescence, but no higher-fluorescence region is present within the SMC domain, indicating complete lack of coacervation. Encapsulating mixture consisted of 7.3 mM DEX (unlabelled: cy5-labeled = 1000:1, molar ratio), 8.6 mM PEG, 2.4 mg/ml PLL (unlabelled: FITC-labeled = 5:1, mass ratio), 0.8 mM ATP, and 15% v/v glycerol in 25 mM citrate-HCl (pH 4). This mixture was combined with an equal volume of feeding aqueous solution containing 600 mM sucrose and 15% v/v glycerol in 25 mM Tris-HCl (pH 9). Scale bar, 50 µm.

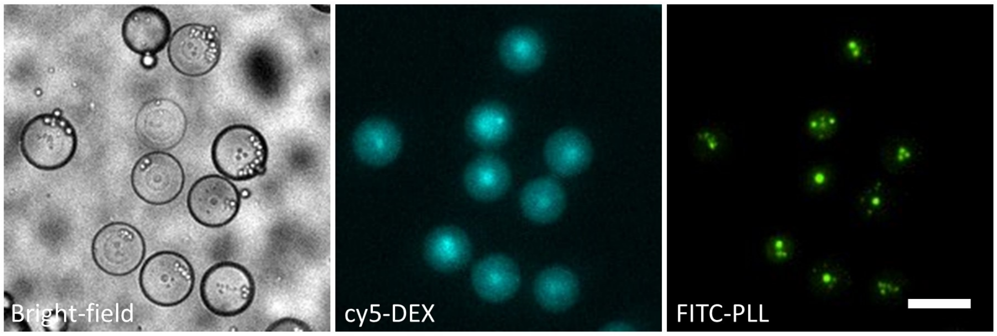


**Figure S4**: **Simultaneous SPS and APS triggers form coacervates within SMC domains.** In case of both triggers, the bright-field image shows a multiphase structure in each double emulsion. The largest domain is an SMC, as confirmed by the DEX fluorescence. The SMC domain in turn shows confined coacervates, as confirmed by the corresponding PLL fluorescence. Encapsulating mixture consisted of 7.3 mM DEX (unlabelled: cy5-labeled = 1000:1, molar ratio), 8.6 mM PEG, 2.4 mg/ml PLL (unlabelled: FITC-labeled = 5:1, mass ratio), 0.8 mM ATP, and 15% v/v glycerol in 15 mM citrate-HCl (pH 4). This mixture was combined with an equal volume of feeding aqueous solution containing 500 mM sucrose and 15% v/v glycerol in 15 mM Tris-HCl (pH 9). Scale bar, 50 µm.

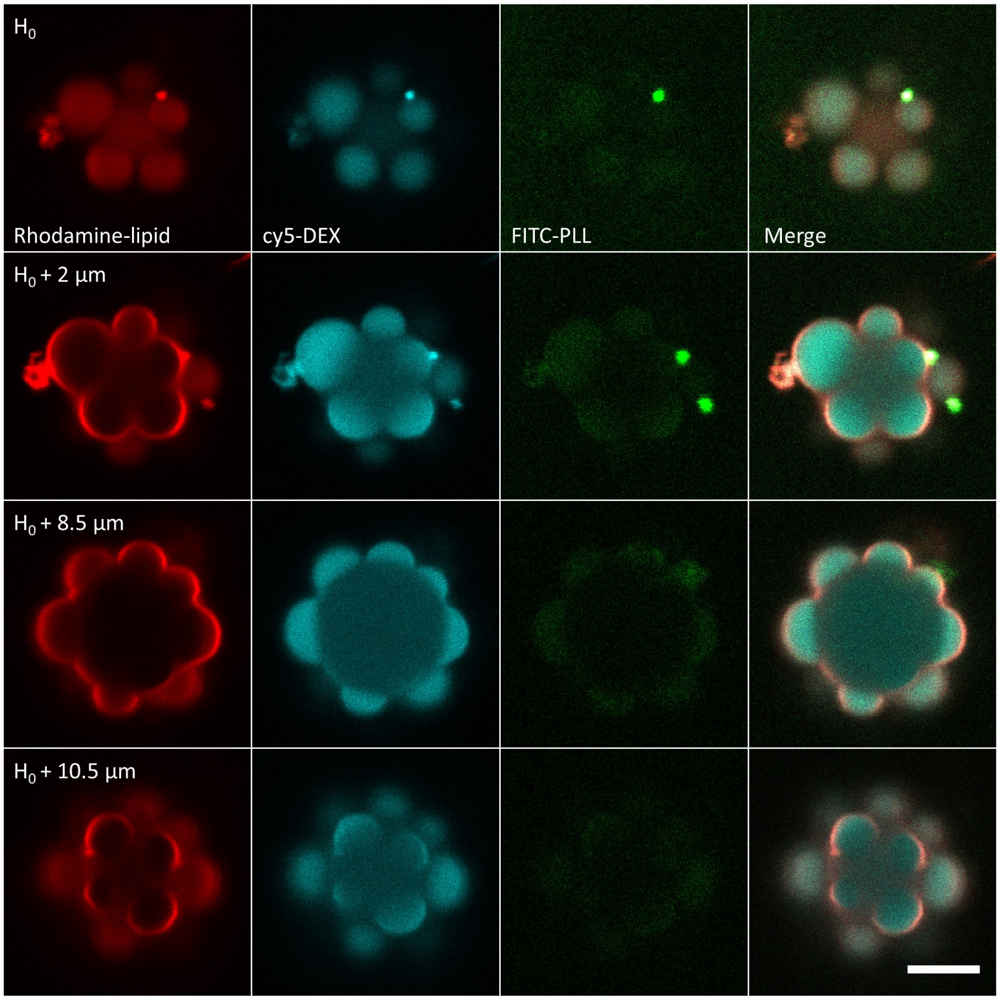


**Figure S5**: **Confocal fluorescence images showing the different planes of a 'flower-shaped' liposome with the SMC 'petals' harboring condensates.** The lipid channel shows the pronounced non-spherical morphology of the liposome. The DEX fluorescence shows the the presence of SMCs that form DEX-rich domains. The PLL fluorescence shows two coacervates confined within two different SMCs. The lipid composition was 99.9 % DOPC + 0.1% Liss Rhodamine PE (molar ratio). Encapsulating mixture consisted of 7.3 mM DEX (unlabelled: cy5-labeled = 1000:1, molar ratio), 8.6 mM PEG, 2.4 mg/ml PLL (unlabelled: FITC-labeled = 5:1, mass ratio), 0.8 mM ATP, and 15% v/v glycerol in 15 mM citrate-HCl (pH 4). This mixture was combined with an equal volume of feeding aqueous solution containing 500 mM sucrose and 15% v/v glycerol in 15 mM Tris-HCl (pH 9). Scale bar, 5 µm.

**Table S1**: **Detailed overview of the compositions used for the inner aqueous (IA), outer aqueous (OA), exit well solution (EX) and feeding aqueous (FA).** Table values denote concentrations of the various constituents used in the experiments. Concentrations are denoted as (i/ii/iii/iv), where the numbers i-iv respectively indicate the concentrations used in i: Fig.2 and Supplementary Fig. 2; ii: Fig. 3 and Supplementary Fig. 3; iii: Fig. 4 and Supplementary Fig. 4b, 5; iv: Supplementary Fig. 4a.

| Compound | IA | OA | EA | FA |
|---|---|---|---|---|
| DEX (mM) | 7.3/7.3/7.3/7.3 | | | |
| Alexa Fluor 647-DEX ($\mu$M) | 7.3/7.3/7.3/7.3 | | | |
| PEG 8k (mM) | 8.6/8.6/8.6/8.6 | | | |
| PLL (mg/ml) | 2/2/2/2 | | | |
| FITC-PLL (mg/ml) | 0.4/0.4/0.4/0.4 | | | |
| ATP (mM) | 0.8/0.8/0.8/0.8 | | | |
| Glycerol (%, v/v) | 15/15/15/15 | 15/15/15/15 | 15/15/15/15 | 15/15/15/15 |
| F68 (%, w/v) | | 5/5/5 | | |
| Sucrose (mM) | | 70/70/70/70 | 70/70/70/70 | 500/600/500/500 |
| Citrate-HCl, pH 4.50 (mM) | 15/0/15/15 | 15/0/15/15 | 15/0/15/15 | 0/0/0/15 |
| Citrate-HCl, pH 4.17 (mM) | 0/25/0/0 | 0/25/0/0 | 0/25/0/0 | |
| Tris-HCl, pH 8.64 (mM) | | | | 15/0/15/0 |
| Tris-HCl, pH 8.95 (mM) | | | | 0/25/0/0 |

**Caption for Movie S1. Restricted motion of coacervates within MSCs, accentuated by an external fluid flow.** We obtained coacervate-in-MSC multi-phase emulsion droplets after mixing the four components under APS and SPS conditions. The coacervates (visualized using FITC-PLL fluorescence and seen as dark green droplets) remain confined within the DEX-rich domains even in presence of an external fluid flow.

**Caption for Movie S2. Two-dimensionally restricted movement of PLL-recruited SMCs on the membrane surface.** The motions of several membrane-bound SMCs (visualized by FITC-PLL fluorescence, in green) are restricted along the membrane surface (visualized by Rhodamine-PE fluorescence, in red).

**Caption for Movie S3. Formation of SPS and APS after exposing the liposome to a hypertonic, high-pH buffer.** After exposing the liposomes with a hypertonic, high-pH buffer, the liposome (visualized by Liss Rhodamine-PE fluorescence, in red) undergoes a volume reduction with the extra lipids forming a pocket as indicated. Over time, DEX-rich domains (visualized by Alexa Fluor 647-DEX fluorescence, in cyan) were formed, underwent coalescence, and ultimlately wetted the membrane. These domains recruited PLL molecules and induced coacervation (visualized by FITC-PLL fluorescence, in green) at the membrane.

**Caption for Movie S4. 3D projection of the final morphology of a liposome 20 minutes after the APS and SPS were triggered.** Multiple 'petals' (visualized by Alexa Fluor 647-DEX fluorescence, in cyan) could be seen, drastically restructuring the liposome into a 'flower' shape. Some 'petals' harbored the formed coacervates (visualized by FITC-PLL fluorescence, in green) as buds on membrane. The lipid membrane (visualized by Liss Rhodamine-PE fluorescence, in red) encapsulated SMC and buds.

**Caption for Movie S5. Confocal fluorescence *z*-stack showing a liposome at different planes after the formation of APS in SMCs.** From bottom to up, multiple 'petals' (visualized by Alexa Fluor 647-DEX fluorescence, in cyan) could be seen, drastically restructuring the liposome into a 'flower' shape. The liposomes show different morphologies in different planes due to the random distribution of the ‘petals’. Some petals harbored the formed coacervate buds (visualized by FITC-PLL fluorescence, in green), and the coacervates are also covered by lipid membrane (visualized by Liss Rhodamine-PE fluorescence, in red).

**Caption for Movie S6. Corralled diffusion of coacervates in SMCs at the liposome membrane.** The lipid membrane is visualized by Liss Rhodamine-PE fluorescence, in red. Due to the separation of the DEX-rich domains (visualized by Alexa Fluor 647-DEX fluorescence, in cyan), the coacervates (visualized by FITC-PLL fluorescence, in green) remained isolated and could not come in contact with coacervates residing in other petals. The coacervate trajectories in each ’petal’ are random but restricted within corresponding SMCs.

**Caption for Data S1.** Raw data underlying Figures 1-4 in the main article and Supplementary Figure 1 in Supplementary information.

**Caption for Data S2.** MATLAB scripts used for image processing and related calculations. All codes used in the figures are listed separately as individual files.
